# Nitrate-mediated luminal expansion of *Salmonella* Typhimurium is dependent on the ER stress protein CHOP

**DOI:** 10.1101/2023.11.03.565559

**Authors:** Lydia A. Sweet, Sharon K. Kuss-Duerkop, Mariana X. Byndloss, A. Marijke Keestra-Gounder

**Author notes:** Correspondence: Marijke Keestra-Gounder, PhD, Department of Immunology and Microbiology, University of Colorado School of Medicine, Aurora, CO 80045.

## Abstract

*Salmonella* Typhimurium is an enteric pathogen that employs a variety of mechanisms to exploit inflammation resulting in expansion in the intestinal tract, but host factors that contribute to or counteract the luminal expansion are not well-defined. Endoplasmic reticulum (ER) stress induces inflammation and plays an important role in the pathogenesis of infectious diseases. However, little is known about the contribution of ER stress-induced inflammation during *Salmonella* pathogenesis. Here, we demonstrate that the ER stress markers *Hspa5* and *Xbp1* are induced in the colon of *S*. Typhimurium infected mice, but the pro-apoptotic transcription factor *Ddit3,* that encodes for the protein CHOP, is significantly downregulated. *S*. Typhimurium-infected mice deficient for CHOP displayed a significant decrease in inflammation, colonization, dissemination, and pathology compared to littermate control mice. Preceding the differences in *S*. Typhimurium colonization, a significant decrease in *Nos2* gene and iNOS protein expression was observed. Deletion of *Chop* decreased the bioavailability of nitrate in the colon leading to reduced fitness advantage of wild type *S*. Typhimurium over a *napA narZ narG* mutant strain (deficient in nitrate respiration). CD11b+ myeloid cells, but not intestinal epithelial cells, produced iNOS resulting in nitrate bioavailability for *S*. Typhimurium to expand in the intestinal tract in a CHOP-dependent manner. Altogether our work demonstrates that the host protein CHOP facilitates iNOS expression in CD11b+ cells thereby contributing to luminal expansion of *S*. Typhimurium via nitrate respiration.

**Author Summary:** *Salmonella* Typhimurium is a gastroenteric bacterium that replicates to large numbers within the gastrointestinal (GI) tract allowing for efficient host-to-host transmission. One strategy that allows *Salmonella* to expand in the GI tract is via nitrate respiration that is generated during *Salmonella* infections. Our results presented here provide more insight into the role of the host protein CHOP in the production of nitrate and the subsequent growth of *Salmonella* in the GI tract. CHOP expression is regulated within the unfolded protein response (UPR), an adaptive response pathway that is activated when cells are undergoing endoplasmic reticulum (ER) stress. ER stress has been implicated in several infectious and inflammatory diseases; however, little is known about the contribution of ER stress and the UPR during *Salmonella* infections. Here, we demonstrate that *Chop* is downregulated in mice infected with *S*. Typhimurium, and that mice deficient for CHOP have reduced bacterial numbers in the colon, suggesting that downregulation of *Chop* is a host response to resist intestinal colonization by *Salmonella*. Our results further show that CHOP contributes to increased expression of iNOS, responsible for nitrate production, thereby increasing the bioavailability of nitrate that allows for *Salmonella* growth. Altogether, our research provides a better understanding of the contribution of the ER stress protein CHOP in intestinal health and disease.

## Introduction

*Salmonella enterica* serovar Typhimurium (*S*. Typhimurium) is a Gram-negative enteric bacterial pathogen that actively induces intestinal inflammation which allows *S*. Typhimurium to outcompete the host microbiota and expand within the gastrointestinal (GI) tract (1, 2). The main virulence factors required for *Salmonella* to induce intestinal inflammation are two type III secretions systems (T3SS) (1–4). The T3SS-1 allows *S*. Typhimurium to invade intestinal epithelial cells (IECs) resulting in the production of cytokines/chemokines, including KC and CCL2, thereby attracting inflammatory phagocytes (macrophages, monocytes, neutrophils and dendritic cells) to the site of infection (3). The T3SS-2 is required for survival and replication within macrophages (5). In addition to the T3SSs, *Salmonella* induces inflammatory responses via Pathogen-Associated Molecular Patterns (PAMPs) that are detected by Pattern Recognition Receptors (PRRs) (6, 7). Early innate immune responses during *S*. Typhimurium infection include the production of an array of chemokines and cytokines such as KC, TNFα, IL-1β and IL-23 (8–10). Intestinal epithelial cells are the predominant producers of neutrophil chemo-attractant KC during *S*. Typhimurium infection (9, 11, 12). TNFα and IL-1β are mainly produced by accumulated inflammatory monocytes and IL-23, produced by dendritic cells, is a crucial cytokine in the development and maintenance of the pro-inflammatory Th17 response (9, 13). In addition to the induction of chemokines and cytokines, *S*. Typhimurium also causes robust expression of *Nos2*. *Nos2* is the gene encoding for inducible nitric oxide (NO) synthase (iNOS) that catalyzes the production of nitric oxide from L-arginine. Nitric oxide radicals react with superoxide radicals which leads to the generation of nitrate (NO ^-^). Nitrate enhances the growth of *S*. Typhimurium via nitrate respiration (14, 15). It was demonstrated that different cells in the intestinal tract produce nitrate, but interestingly, only the nitrate produced by phagocytic infiltrates is utilized by *S*. Typhimurium (16, 17).

Upon infection, the intestinal tract must cope with diverse cellular stresses and as a response, cells activate mechanisms to support cellular functions to adapt to changing environmental conditions. Activation of the unfolded protein response (UPR) is one such mechanism that is induced upon endoplasmic reticulum (ER) stress (18). The ER is a highly dynamic organelle that exerts a major role in coordinating signaling pathways to ensure cellular homeostasis. The ER is the site of synthesis and folding of proteins; however, under different stressful pathological and physiological conditions, the ER is unable to maintain homeostasis and activates the UPR (18). Three transmembrane receptors, ATF6, PERK and IRE1α, are activated and regulate biological processes such as inhibition of protein translation, autophagy, and inflammation to restore cellular homeostasis. Under homeostatic conditions, the chaperone BiP, encoded by the *Hspa5* gene, is bound to these receptors, thereby preventing their activation. Perturbation of the ER triggers the release of BiP from ATF6, PERK and IRE1α, resulting in dimerization and phosphorylation of these receptors to an active state (18). ATF6, PERK and IRE1α subsequently activate the transcription factors ATF6f, ATF4 and XBP1, respectively, which then bind to ER stress elements (ERSE) that result in the transcription of genes such as *Hspa5*, *Xbp1*, and *Ddit3* (herein referred to as *Chop*), the gene encoding transcription factor CCAAT/enhancer-binding protein homologous protein (CHOP) (18). CHOP is a downstream apoptosis-promoting target of the PERK-ATF4 pathway, resulting in ER stress-induced cell death when the ER stress is severe and/or prolonged (19). Although the role of CHOP has focused mostly on the induction of cell death, more recent studies have reported a role for CHOP in regulating inflammation during infections (20, 21). Infection of myeloid cells with *Chlamydia trachomatis* resulted in increased *Chop* expression and binding of CHOP to the *IL23* promoter (21). Increased expression of *Chop* mRNA and CHOP protein was also detected in trophoblasts infected with *Brucella abortus* (22). Activation of *Chop* during *Listeria monocytogenes* infection was shown to be detrimental to the host leading to increased morbidity and mortality (23). Shiga toxins expressed by enteric pathogens *Shigella dysenteriae* 1 and enterohaemorrhagic *Escherichia coli* (EHEC) increased CHOP expression and induced monocytic cell apoptosis (24). Thus, CHOP has a critical, yet not well defined, role during bacterial infections.

Although a lot is known about the mechanisms of *Salmonella*-induced intestinal inflammation, not much is known about the contribution of ER stress during *Salmonella* infections. Treatment of *Salmonella*-infected mice with the ER stress inhibitor TUDCA resulted in increased colonization of the small intestine because of decreased lysozyme production mediated by the ER stress-induced secretory autophagy pathway in Paneth cells (25). In HeLa cells, however, it was demonstrated that cells pretreated with the ER stress inducer thapsigargin resulted in increased *Salmonella* replication, suggesting that activation of ER stress provides a favorable environment for *Salmonella* (26). Here, we set out to investigate the involvement of ER stress, and more specifically the function of ER stress-induced CHOP, in the *Salmonella*-induced colitis model as well as the contribution of CHOP to inflammation, colon pathology and luminal expansion of S. Typhimurium.

## Results

### Activation of the ER stress response during *S*. Typhimurium infection

To investigate whether the ER stress response is activated in the colon during infection we infected streptomycin pretreated mice with *S*. Typhimurium SL1344 and humanely euthanized the mice at 24, 48 and 72 hr post infection. Expression of the UPR target genes *Xbp1* and *Hspa5* are significantly upregulated in the colon of infected mice at all time points tested, indicating the ER stress response is activated during infection (Figure 1A and B). In contrast, *Chop* is significantly downregulated at day 72h post infection (Figure 1C). These results suggest that ER stress is activated during *Salmonella* infection, but that activation of the three UPR branches may be differentially regulated during intestinal inflammation.

**Figure 1.**
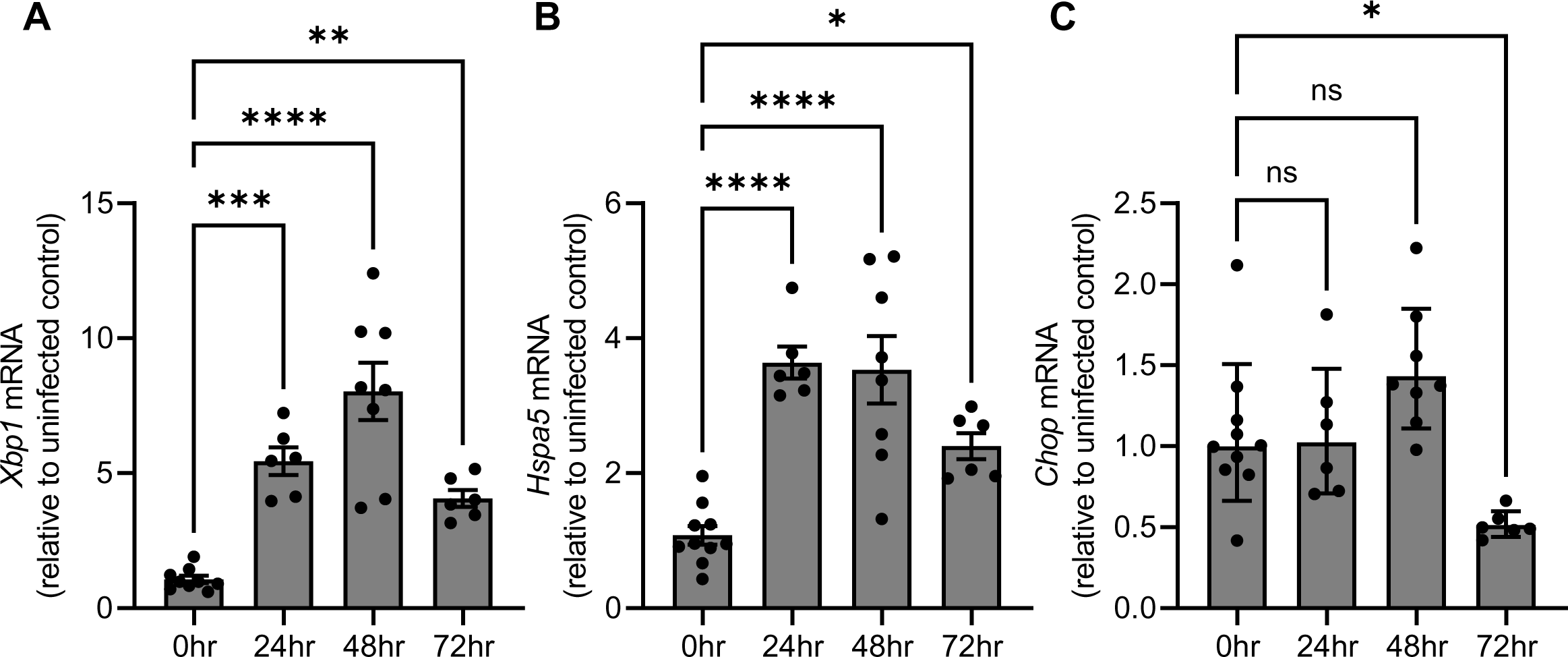
*S.* Typhimurium activates the UPR *in vivo.* RNA was extracted from the colon of S. Typhimurium (SL1344) infected (10^8^ cfu/mouse) and uninfected mice (C57BL/6) and analyzed by qRT-PCR for expression of the ER stress markers *Xbp1* (A), *Hspa5* (B) and *Chop* (C). Data shown as mean ± SEM with 6-10 mice per group. Ordinary one-way ANOVA followed by Dunnett’s multiple comparisons test. p value *<0.05, **<0.01, ***<0.001 and ****<0.0001 using GraphPad PRISM.

### *Chop^-/-^* mice have reduced *S*. Typhimurium colonization, dissemination and tissue pathology

The dissociation between UPR activation and reduced *Chop* expression in *Salmonella*-infected colonic tissue was surprising, since it has been demonstrated previously that other bacterial pathogens induce the expression of *Chop* (21–25). To investigate the role of CHOP during *Salmonella* infections, we infected CHOP deficient mice (*Chop^-/-^*), heterozygous (*Chop^+/-^*) and wildtype (*Chop^+/+^*) control mice with *S*. Typhimurium SL1344. Expression of *Xbp1* and *Hspa5* was not significantly different in *Chop^-/-^* mice compared to the control mice, indicating ER stress is induced in CHOP deficient mice (Figure S1A and B). Importantly, no significant differences in colon colonization and dissemination to the liver were observed when comparing heterozygous (*Chop^+/-^*) and wildtype (*Chop^+/+^*) mice (Figure S1C and D). We therefore continued using heterozygous *Chop^+/-^* littermate control mice in comparison with *Chop^-/-^* mice for consecutive experiments. The *Chop^-/-^* mice, on the other hand, had significantly lower bacterial numbers in the colon and liver 72hr post infection (Figure 2A and B). At 72hr post infection the *Chop^-/-^* mice had significantly reduced histopathology scores and neutrophil counts (Figure 2C-E). There were no significant differences in histopathology scores between the *Chop^-/-^* mice and the littermate control mice at 48hr post infection (Figure S2).

**Figure 2.**
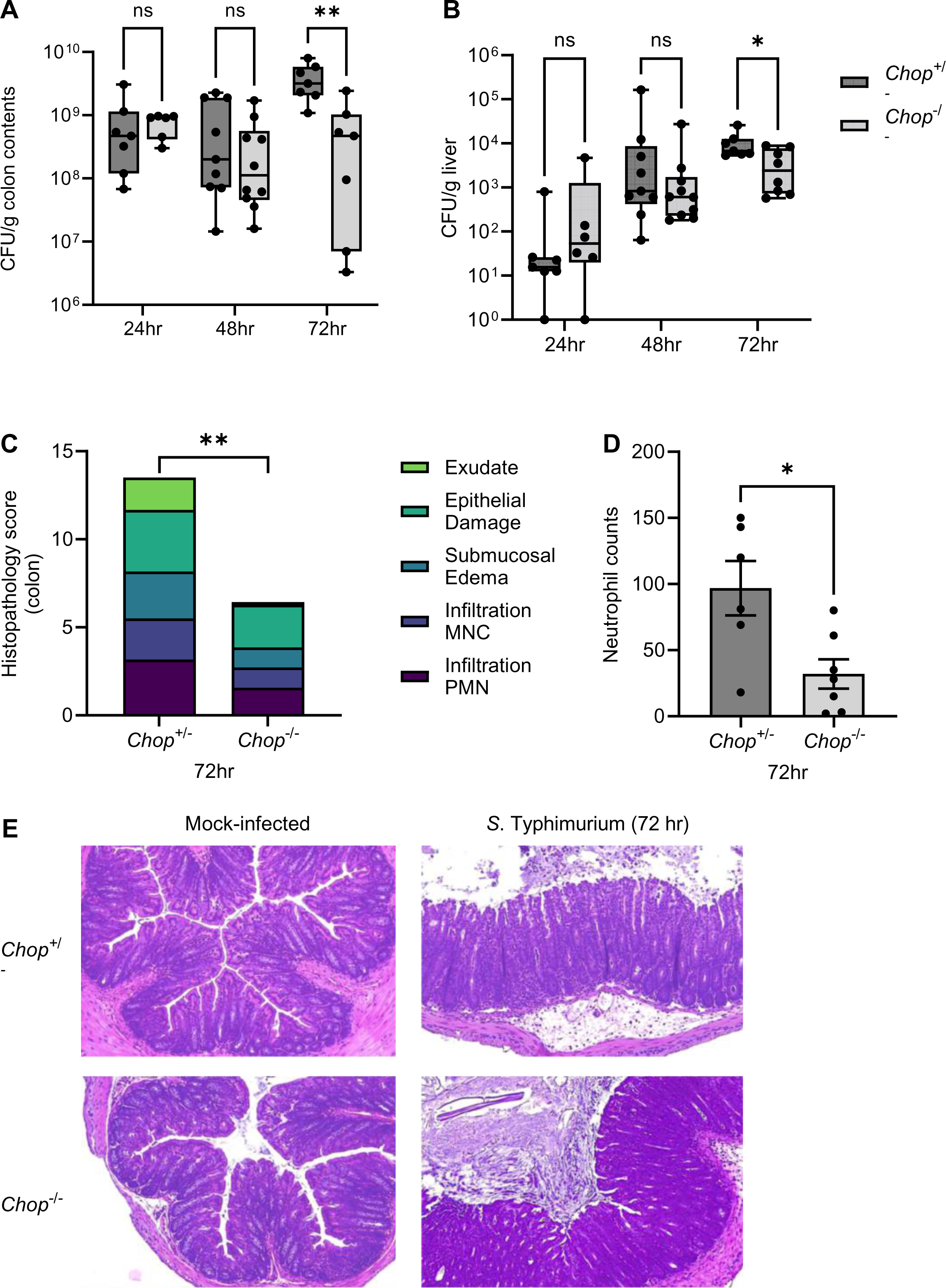
CHOP contributes to colonization, dissemination, and pathology during S. Typhimurium infection. Streptomycin-pretreated *Chop^-/-^* and *Chop^+/-^*littermate control mice were infected with *S*. Typhimurium (SL1344, 10^8^ cfu/mouse) and bacterial numbers were determined in colon contents (A) and liver (B) at 24h, 48h and 72h post infection. (C) Total histopathology scores for colon sections at 72h p.i. (D) Total neutrophil count. (E) Representative images of the colon of *Chop^-/-^* and *Chop^+/-^*mice mock-treated or infected with S. Typhimurium for 72h. Data shown as mean ± SEM or min to max with 6-10 mice per group. A and B; Multiple unpaired t tests. C and D; unpaired t tests. p value *<0.05, **<0.01, ns (not significant) using GraphPad PRISM.

### CHOP contributes to the expression of pro-inflammatory cytokines during *S*. Typhimurium infection

*S*. Typhimurium utilizes inflammation to drive bacterial expansion, suggesting that the reduced bacterial burdens and histopathology at 72h p.i. may be linked to a decrease in the inflammatory response (2) (Figure 2). At 24 hours post-infection we observed decreased expression of *Nos2* and *Tnfa* in the colon of *Chop^-/-^* mice suggesting that CHOP has a role in the induction of inflammation (Fig. 3A and B). We did not observe any differences at basal level in mock-infected *Chop^-/-^* mice and littermate control mice (data not shown). At the later time points, *Nos2* and *Tnfa* expression remained decreased and *Il1b, Kc*, and *Il23* were also reduced (Figure 3). The reduced gene expression at 24 and 48hr post infection was not the result of reduced bacterial numbers in the colon, since the difference in colon colonization was observed only at 72hr post infection (Figure 2A and B).

**Figure 3.**
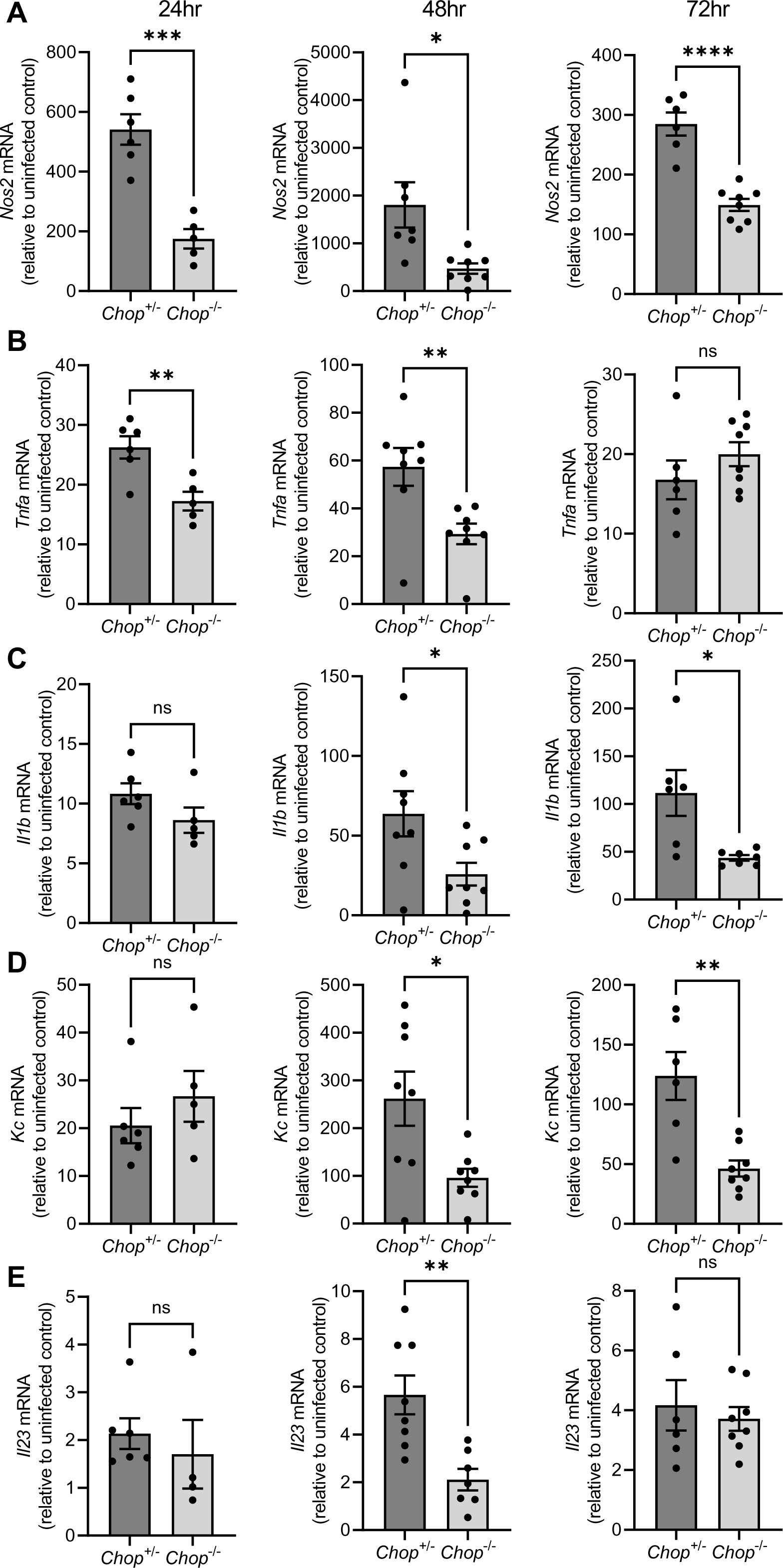
*Chop^-/-^* mice have reduced cytokine expression. RNA from the colon of infected and uninfected *Chop^-/-^* and *Chop^+/-^* littermate control mice was extracted and analyzed by qRT-PCR to determine the levels of *Nos2* (A), *Tnfa* (B), *Il1b* (C), *Kc* (D), and *Il23* (E) at 24h, 48h and 72h post infection. Data shown as mean ± SEM with 5-8 mice per group. Unpaired t tests. p value *<0.05, **<0.01, ***<0.001 and ****<0.0001, ns (not significant) using GraphPad PRISM.

### *Chop^-/-^* mice have reduced iNOS expression leading to decreased colon colonization

*Nos2* encodes the inducible nitric oxide synthase (iNOS), which is present at lower levels in the colon of *Chop^-/-^* mice 48hr after infection with *S*. Typhimurium (Figure S3A). The reduced expression of *Nos2* mRNA and iNOS protein is of particular interest, because iNOS has been implicated in enhanced nitrate production in the intestinal lumen, thereby contributing to *S*. Typhimurium replication via nitrate respiration (14, 15). To investigate whether deletion of *Chop* would decrease the bioavailability of nitrate in the colon, *Chop^-/-^* and heterozygous control mice were infected with a 1:1 mixture of wild type SL1344 and a nitrate respiration-deficient strain (*napAnarGnarZ*) (15). At 48hr and 72hr post infection mice were sacrificed and CFUs in the colon contents enumerated. The fitness advantage conferred to the wild type SL1344 by nitrate respiration was significantly diminished in the *Chop^-/-^* mice at 72hr, but not at 48hr post infection (Figure 4 and Figure S3B). These results suggest a role for CHOP in the production of nitrate resulting in the growth benefit for *S*. Typhimurium via nitrate respiration.

**Figure 4.**
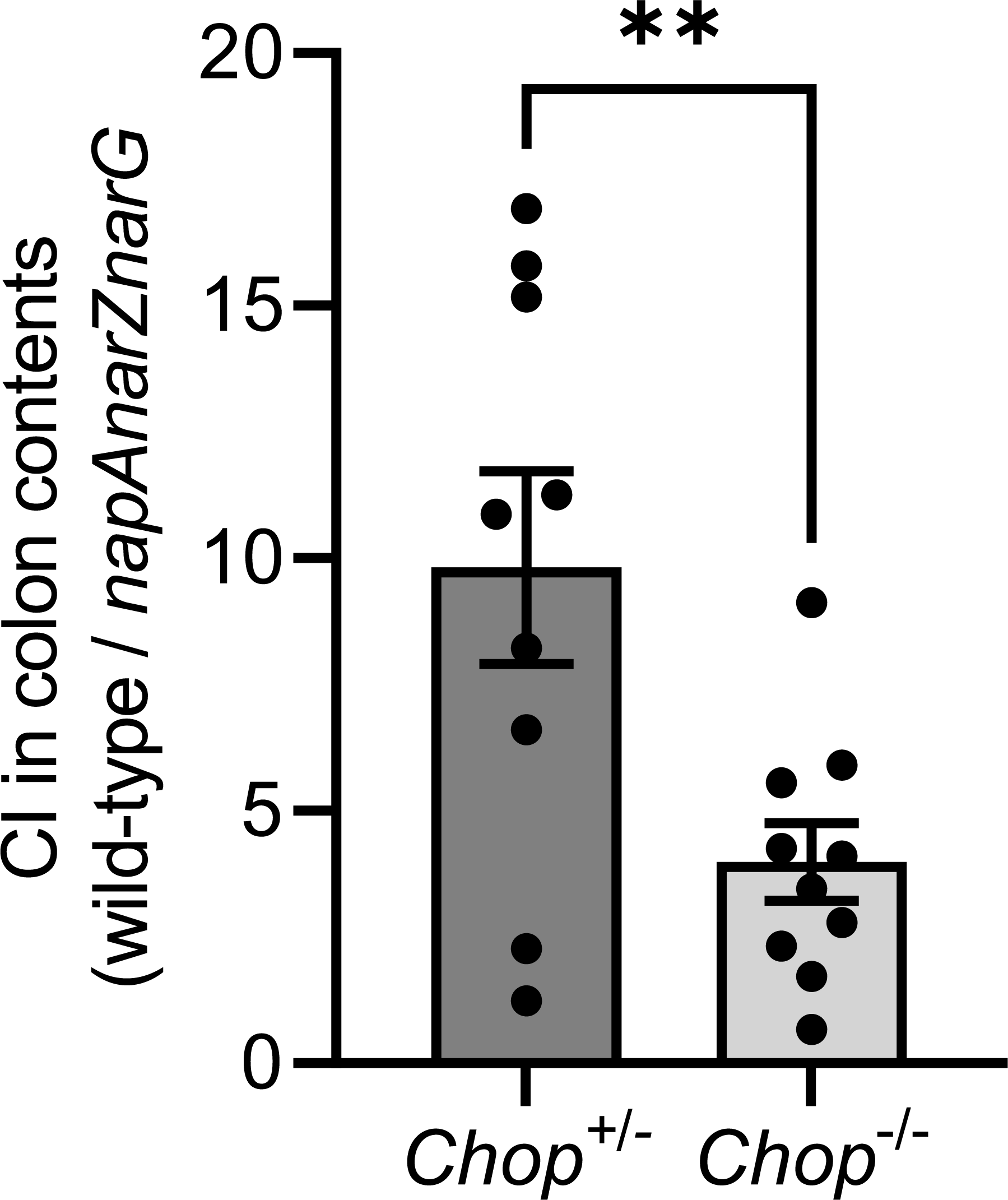
CHOP contributes to nitrate production resulting in a growth benefit for *S*. Typhimurium. Streptomycin-pretreated *Chop^-/-^* and *Chop^+/-^* littermate control mice were infected with a 1:1 mixture of wildtype *S*. Typhimurium SL1344 and *ΔnapAnarZnarG* (10^8^ cfu/mouse). The competitive index (CI) in the colon contents was determined 72h post infection. Data shown as mean ± SEM with 9-10 mice per group. Unpaired t test. p value **<0.01 using GraphPad PRISM.

### CHOP expression in intestinal epithelial cells does not contribute to *S*. Typhimurium pathogenesis

Previous studies with mouse models of T-cell-mediated and bacteria-driven colitis have demonstrated that both CHOP mRNA and protein expression is downregulated in intestinal epithelial cells (27). To investigate whether CHOP expression in IECs results in increased *Salmonella* burden we infected *Chop*^ΔIEC^ mice and littermate control mice (*Chop^flox^*) with *S.* Typhimurium. At 48hr and 72hr post infection we did not observe any significant differences in colon colonization or dissemination to the liver (Figure 5A and B, and Figure S4A and B). Moreover, no differences were observed in histopathology scores or neutrophil counts (Figure 5C and D, and Figure S4C and D), suggesting that CHOP expression in IECs does not contribute to *Salmonella* expansion or pathology in the intestinal tract. Additionally, we did not observe significant differences in gene expression of *Kc*, *Il1b*, *Tnfa* and *Il23* (Figure S4 and S5). More importantly, no differences were observed in the expression of *Nos2* (Figure 5E), indicating that CHOP in IECs does not contribute to nitrate production required for *Salmonella* growth in the intestinal lumen.

**Figure 5.**
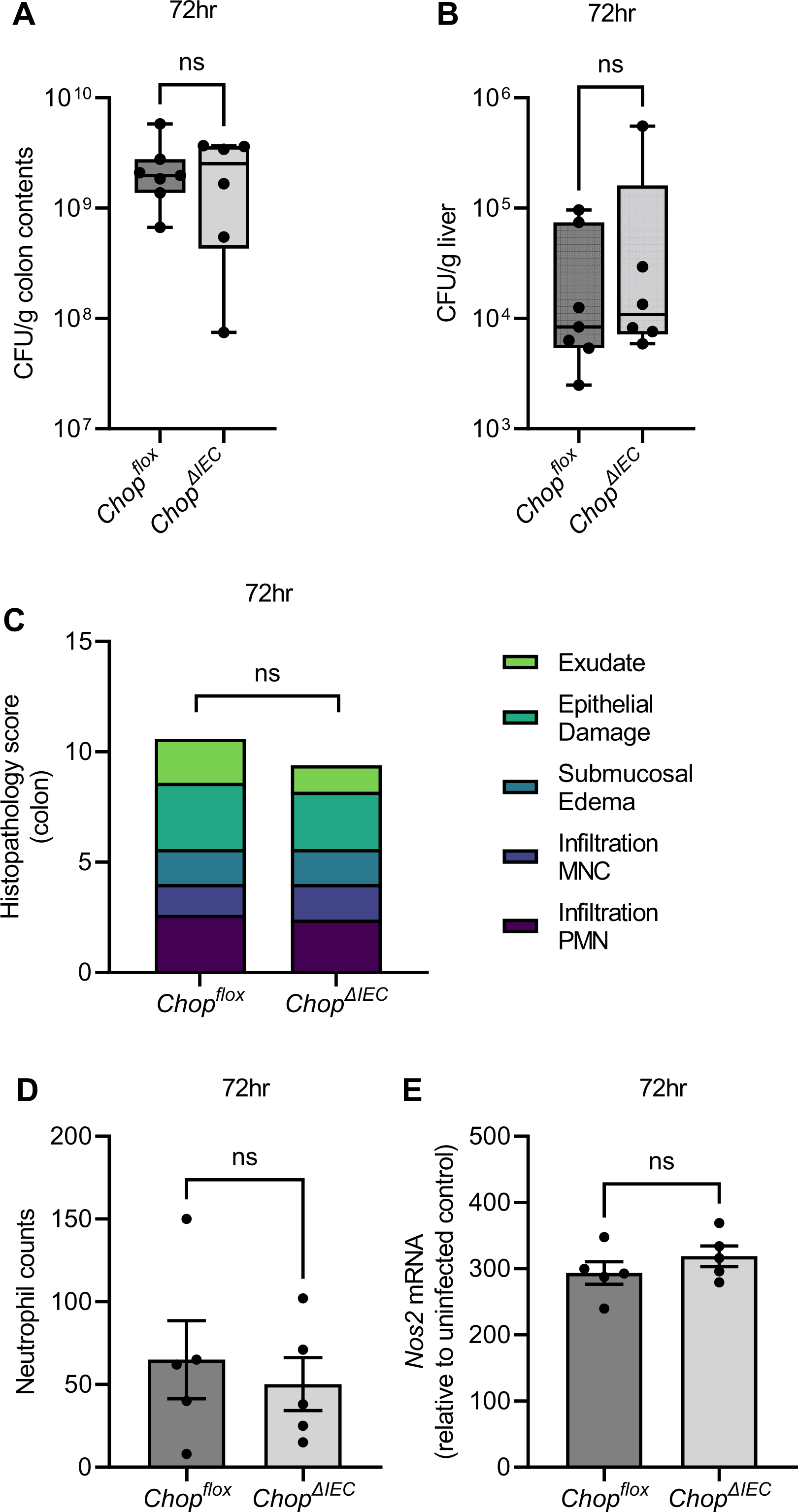
CHOP expression in IECs does not contribute to resistance to S. Typhimurium infections. Streptomycin-pretreated *Chop^ΔIEC^* and *Chop^flox^* littermate control mice were infected with *S*. Typhimurium (SL1344, 10^8^ cfu/mouse) and cfus were determined in colon contents (A) and liver (B) at 72hpi. (C) Total histopathology scores and (D) neutrophil counts in the colon at 72hpi. (E) *Nos2* mRNA levels in the colon. Data shown as mean ± SEM or min to max with 5-7 mice per group. (A and B) Mann-Whitney U test, (C-E) unpaired t tests, ns (not significant).

### iNOS expression in CD11b+ colonic cells is dependent on CHOP

It was demonstrated that inflammatory phagocytes produce nitrate which is utilized by S. Typhimurium (16, 17). To investigate whether CHOP expression in myeloid cells contributes to the production of nitrate, flow cytometric analysis was performed on live colonic cells for the expression of iNOS (Figure 6A). Isolated live cells were gated for a population that was positive for the leucocyte marker CD45 and positive for CD11b, the marker for inflammatory phagocytes (i.e. monocytes/macrophages, neutrophils). There were no significant differences in the percentage of CD45+/CD11b+/iNOS-cells isolated from *S.* Typhimurium infected *Chop^-/-^* mice compared to littermate control mice (Figure 6B), suggesting CHOP expression is not required for recruitment of CD11b+ cells. Infected *Chop^-/-^* mice, however, have a lower percentage of live CD45+/CD11b+ cells that were iNOS+ compared to infected littermate control mice, indicating that CHOP expression in CD11b+ cells contribute to iNOS production (Figure 6C). Consistent with this finding, flow cytometric analysis disclosed that the total number of CD45+/CD11b+/iNOS-cells from the colon of infected *Chop^-/-^* and littermate control mice was not significantly different, but *Chop^-/-^* mice have significantly lower numbers of CD45+/CD11b+/iNOS+ cells (Figure 6D and E). Overall, our data demonstrate that CHOP contributes to iNOS expression in CD45+/CD11b+ colonic cells, resulting in increased nitrate availability that *S*. Typhimurium utilizes for respiration during infection.

**Figure 6.**
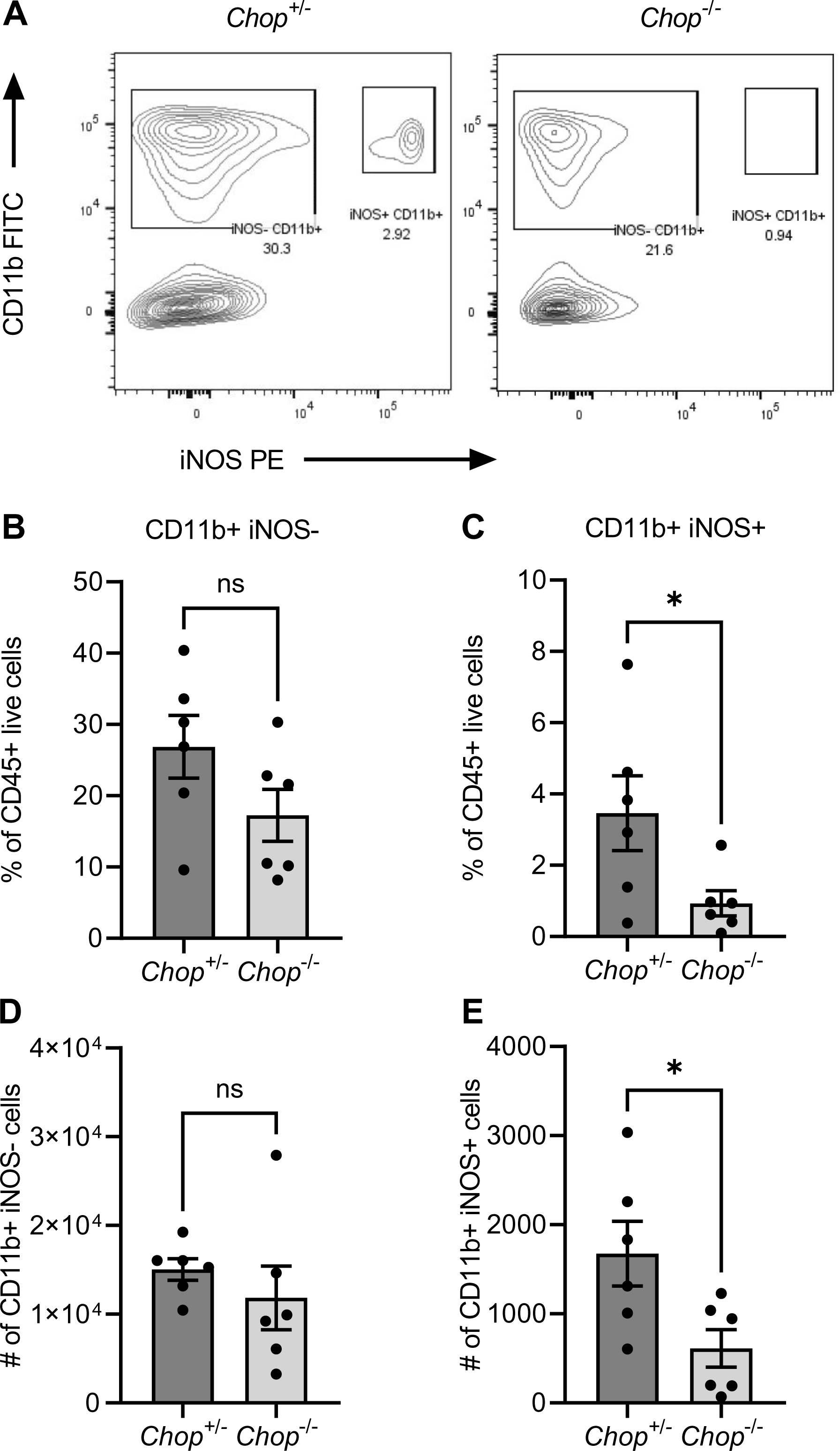
CHOP deficient CD45+/CD11b+ colonic cells produce less iNOS. Representative flow plots of staining for CD11b and iNOS on cells isolated from colonic tissue of *S*. Typhimurium infected mice at 72h p.i. (A). Percent of CD45+ live cells that are CD11b+/iNOS-(B) and CD11b+/iNOS+ (C). Quantification of number of CD11b+/iNOS-cells (D) and CD11b+/iNOS+ cells (E). Data shown as mean ± SEM with 6 mice per group. Unpaired t tests. p value *<0.05, ns (not significant) using GraphPad PRISM.

## Discussion

ER stress and activation of the UPR play major roles in the pathology and resolution of bacterial infections (24, 28–30). Our work expands on the role of ER stress during *S*. Typhimurium infection and indicates that CHOP promotes luminal expansion in the colon via nitrate respiration derived from CD45+CD11b+ cells. Bel *et al* demonstrated that *S*. Typhimurium infection increased CHOP expression in the small intestine, contrary to our finding of *Chop* downregulation in colonic tissues. Activation of ER stress by *S*. Typhimurium in Paneth cells in the mouse small intestine resulted in activation of the secretory autophagy pathway leading to increased lysozyme secretion in the lumen and increased lysozyme mediated killing (25). Treating mice with tauroursodeoxycholic acid (TUDCA), thereby inhibiting the ER stress response, and thus CHOP expression, resulted in increased bacterial numbers in the small intestinal luminal contents and increased dissemination to liver and spleen. TUDCA attenuates ER stress by inhibiting the dissociation of BiP from the receptors and therefore may not only inhibit CHOP expression but also the IRE1α and ATF6 pathways, which could contribute to the increased bacterial numbers (31). CHOP protein and *Chop* mRNA is downregulated in colonic tissues derived from mouse models of T-cell-mediated and bacteria-driven colitis (27) and from colonic tissues from ulcerative colitis (UC) patients (32). In DSS- and TNBS-induced colitis, however, CHOP protein and *Chop* mRNA was increased (33). Why CHOP is differentially regulated in a variety of colitis and infection models requires further investigation. Downregulation of CHOP might be a protective mechanism of the host to limit inflammation and/or cell death, whereas upregulation of CHOP may increase inflammation and cell death that could help the host fight off infections. Since *Salmonella* benefits from inflammation, one could argue that the reduced expression of *Chop* in colonic tissue is a protective host response to prevent luminal expansion.

Specific deletion of *Chop* from IEC did not contribute to *S*. Typhimurium-induced colitis (Figure 5). We observed no differences in pathology, inflammatory responses, or *Salmonella* burdens in colon and liver in *Chop*^ΔIEC^ mice compared to littermate control mice. In contrast, mice with IEC-specific overexpression of CHOP were more susceptible to DSS-induced colitis, suggesting an important role for CHOP in intestinal epithelial cells during chemically-induced gut inflammation (27). One of the hallmarks of disease in these mice was an increased number of apoptotic IECs compared to wildtype control mice, indicating an important role for CHOP-mediated cell death. Although we did not specifically test for increased IEC apoptosis, *Chop*^ΔIEC^ mice did not display increased histopathological scores which includes epithelial damage, suggesting that *S*. Typhimurium-induced IEC apoptosis occurs via CHOP-independent mechanisms (34). Our findings, however, do not exclude a role for CHOP in the induction of IEC apoptosis during *Salmonella* infections, as deletion of one cell death pathway may be compensated by other cell death pathways. It has recently been demonstrated that there is an extensive cross-talk between initiators and effectors of distinct cell death pathways, and that loss of a single pathway does not significantly contribute to control of *Salmonella*, but that combined deletion of multiple cell death pathways causes loss of bacterial control in mice (34).

Our results revealed that CHOP contributes to the upregulation of several pro-inflammatory mediators, including *Nos2, Kc*, *Il1b*, *Tnfa* and *Il23*. At day 1 and day 2 post infection we observed significantly reduced expression of these cytokines, but no differences in bacterial numbers in colon contents and liver, indicating the reduced gene expression was not a consequence of reduced bacterial numbers. These differences were only observed in infected animals, as there were no significant differences in background expression of these cytokines in uninfected *Chop^-/-^* mice and littermate control mice (data not shown). Since activated macrophages/monocytic infiltrates are capable of secreting KC, IL-1β, TNFα, IL-23 and expressing iNOS, and we excluded a role for IECs, we hypothesized that CHOP plays a role in macrophages by regulating the expression of pro-inflammatory cytokines. Our data demonstrates that CD11b+ cells isolated from colonic tissue of *Chop^-/-^* infected mice have reduced iNOS expression, suggesting that CD11b+ cells, which include macrophages/monocytic infiltrates, contribute to the luminal expansion of *S*. Typhimurium at day 3 post infection. Indeed, the role of monocyte-derived nitrate production in *S*. Typhimurium infection was previously demonstrated. McLaughlin *et al.* showed that mice lacking CCR2, which are unable to recruit monocytes to the intestine, have reduced *Nos2* and iNOS expression and reduced nitrate in the cecal lumen (16). More recently Liou *et al* demonstrated that *S*. Typhimurium does not respire nitrate derived from IEC but rather from inflammatory phagocytes (17). Here, we demonstrate that CHOP contributes to nitrate production and the subsequent replication of *S*. Typhimurium. Furthermore, nitrate bioavailability in the colon of *S*. Typhimurium infected mice is not regulated by CHOP expressed in IEC, suggesting that IEC-derived nitrate does not contribute to nitrate utilization by *S*. Typhimurium which is consistent with the findings from Liou *et al*. Interestingly, intestinal inflammation alters the colonic microbiota, supporting the growth of the Enterobacteriaceae family, which includes *Escherichia coli* (35). Winter *et al.* have demonstrated that one of the underlying mechanisms of growth of *E. coli* in the inflamed GI tract is via nitrate respiration (36), which was later confirmed to be derived from IECs and regulated by Peroxisome proliferator-activated receptor gamma (PPAR-γ) (17, 37). Although we demonstrate that CHOP expression in IECs does not contribute to nitrate respiration by *S*. Typhimurium, this does not exclude the possibility that CHOP does regulate iNOS expression in IECs and that CHOP-mediated epithelial-derived nitrate is utilized by *E. coli*. This could explain a possible mechanism of how CHOP expression in IEC contributes to disease severity in IBD patients as well as in DSS- and TNBS-induced colitis models, and why a bloom of *E. coli*, and not *S*. Typhimurium, is associated with IBD (27, 35, 36, 38).

How CHOP controls the expression of iNOS and a variety of cytokines remains to be elucidated. CHOP is a transcription factor and thus might be able to directly influence the transcription of cytokines (21). Alternatively, CHOP may control the expression of pro-inflammatory mediators indirectly by altering the polarity of macrophages (39). Classically activated macrophages (CAM or M1 macrophages) produce excessive quantities of the pro-inflammatory cytokines IL-1, TNFα, and IL-23 as well as increased iNOS expression, whereas alternatively activated macrophages (AAM or M2 macrophages) have characteristics of attenuating inflammation (40). It was demonstrated that CHOP plays a role in polarization of adipose tissue macrophages. High fat diet (HFD) feeding of *Chop^-/-^* mice skewed the recruitment of infiltrating macrophages more towards a M2 phenotype compared to wild type control mice on a HFD (39). This raises the intriguing thought that downregulation of CHOP might not be a protective host response but might be actively controlled by *Salmonella* to increase M2 macrophages which are metabolically different than M1 macrophages and more favorable for *Salmonella* for long-term persistence (41, 42).

The polarization of macrophages might be mediated via CHOP repression of PPAR-γ activation. In IECs it has been demonstrated that ER stress-induced CHOP can act as a repressor of PPAR-γ by sequestering the transcription factor C/EBPβ to prevent it from binding to the promoter region of PPAR-γ, resulting in an increased inflammatory response (43). PPAR-γ activation suppresses the immune state of macrophages by repressing transcription of inflammatory mediators including *Nos2*, *Tnfa*, and *IL1b*. Therefore, in the absence of CHOP, increased PPAR-γ signaling allows for reduced cytokine and *Nos2* expression and subsequently reduced levels of nitrate that can be utilized by Enterobacteriaceae for anaerobic respiration (37).

In summary, our results expand on the role of CHOP in the induction of intestinal inflammation during *S*. Typhimurium infection which increases bacterial expansion through increased nitrate bioavailability. Although expression of the ER target genes *Hspa5* and *Xbp1* are increased, *Chop* mRNA expression is reduced in colonic tissues during *S.* Typhimurium infection, suggesting that downregulation of *Chop* might be regarded as a protective host response mechanism, since *Chop^-/-^*mice have reduced histopathology and *S*. Typhimurium growth. Overall, we demonstrate that the host factor CHOP drives the production of nitrate from CD11b+ cells, which is utilized by *Salmonella* to expand in the intestinal tract.

## Materials and Methods

### Mice

*Chop*^-/-^ (B6.129S(Cg)-Ddit3^tm2.1Dron^/J, strain 005530) mice and wild-type C57BL/6 mice were purchased from Jackson laboratory. *Chop*^-/-^ mice were bred with wildtype mice to generate *Chop*^+/-^ mice and then *Chop*^+/-^ mice were bred with *Chop*^-/-^ mice to get *Chop*^+/-^ and *Chop*^-/-^ littermates for experiments. *Chop^fl/fl^* (B6.Cg-Ddit3^tm1.1Irt^/J, strain 030816) and Villin-Cre (B6.Cg-Tg(Vil1-cre)997Gum/J, strain 004586) mice were purchased from Jackson laboratory. *Chop^fl/fl^* and Villin-Cre mice were bred together to generate *Chop^fl/fl^* Cre- and *Chop^fl/fl^* Cre+ littermate mice. All mice were genotyped using protocols and primers from The Jackson Laboratory.

### *Salmonella* Typhimurium Infection

Male and female mice (6-10 weeks of age) were pre-treated with 20 mg of streptomycin in 100 μL of water by oral gavage 24 hours before infection. Mice were infected with 10^8^ CFU SL1344 in 100 μL LB broth or mock infected with 100 μL LB broth via oral gavage. Mice were weighed daily and sacrificed at 24-, 48-, or 72-hours post-infection. At the time of sacrifice liver and colon contents were collected for bacterial enumeration. Samples for enumeration were homogenized, serially diluted, and plated on plates containing the appropriate antibiotics. Portions of the colon tissue were snap frozen in liquid nitrogen for RNA and protein extraction. Sections of colon tissue were also collected for histology. For competitive index experiments, mice were infected with 5×10^7^ of *S*. Typhimurium SL1344 Kan^r^ and 5×10^7^ of *S*. Typhimurium *narG narZ napA* Kan^r^ Carb^r^ in 100 μL LB broth. Colon contents were collected for plating and antibiotic selection was used to determine the colonization level for each strain. All mouse experiments were approved by the Institutional Animal Care and Use Committees at the University of Colorado Anschutz Medical Campus.

### Quantitative Reverse Transcription-PCR

RNA was isolated from colon tissue using TRI Reagent (Molecular Research Center) according to the manufacturer’s instructions. Reverse transcription was performed using 1 μg of DNase-treated RNA (TURBO DNA-free Kit) with TaqMan Reverse Transcription Reagents (Applied Biosystems). Real-time PCR was performed using SYBR Green PCR Master Mix (Applied Biosystems) and the Quantstudio 7 Flex real-time PCR system (Applied Biosystems). Fold change in mRNA levels was calculated using the delta-delta comparative threshold cycle (Ct) method. All targets were normalized to expression levels of *Gapdh* (Table 2. Primer sequences used for qRT-PCR).

### Western Blot Analysis

For protein extraction from tissue samples, tissue pieces were snap frozen in liquid nitrogen at the time of harvest. RIPA buffer (10 mM Tris HCl, 1 mM EDTA, 1% Triton X-100, 0.1% Sodium deoxycholate, 0.1% SDS, 140 mM Sodium chloride, 1mM Phenylmethylsulfonyl fluoride, protease inhibitor cocktail (Roche)) was added to tissue samples and samples were homogenized using a bead beater and put on ice to lyse for 1 hour. The tissue lysate was centrifuged at 13000 rpm for 15 minutes at 4°C and supernatants containing the proteins were transferred to fresh tubes. Protein concentration was measured using the Pierce BCA protein assay kit according to the manufacturer’s instructions. 30 μg of each sample was run on a 10% SDS-PAGE gel for 1 hour at 150 volts. Proteins were transferred to PVDF membrane for 90 minutes at 90 volts. Membranes were then blocked for 1 hour with 5% milk in TBST at RT. Membranes were incubated with anti-iNOS (1:500) and anti-a/b Tubulin (1:5,000) (Cell Signaling Technologies) overnight at 4°C. Secondary anti-rat antibody was used at 1:10,000 for 1 hour at RT. Membranes were developed using ECL Clarity (Bio-Rad) for 4 minutes and then imaged using a G:BOX (Syngene).

### Histopathology

Histological samples from mouse experiments were collected in 10% buffered formalin until the time they were embedded, sliced, mounted, and H&E stained by the histology core in Dr. Rubin Tuder’s laboratory at the University of Colorado Anschutz Medical Campus. All scoring was done blinded by a veterinary pathologist (Dr. Mariana X. Byndloss at Vanderbilt University). The following pathological changes were scored: (i) neutrophil infiltration, (ii) infiltration by mononuclear cells, (iii) submucosal edema, (iv) epithelial damage, and (v) inflammatory exudate. The pathological changes were scored on a scale from 0 to 4 as follows: 0, no changes; 1, detectable; 2, mild; 3, moderate; 4, severe. Neutrophil counts were determined by high-magnification (x400) microscopy, and numbers were averaged from 10 microscopic fields for each animal.

### Single Cell Isolation

After animal sacrifice, the colon was removed and put in ice cold RPMI. Colons were washed with ice cold PBS to remove any contents. The colons were then cut longitudinally and placed in PBS containing 7.5 mM HEPES and 1 mM EDTA (IEL buffer) and shaken at 37°C for 10 minutes. Tissues were then filtered through a 70 μm filter (Fisher Scientific) and put in fresh IEL buffer and shaken for an additional 10 minutes. The samples were filtered again and minced with scissors. Tissues were then digested in RPMI supplemented with 10% FBS, 0.25% β-mercaptoethanol, 3.75 mM HEPES, and 0.54 mg/mL type IV collagenase (Worthington) for 30 minutes at 37°C. Digested tissue was then forced through a 70 μm filter and single cells were counted using a C20 Automated Cell Counter (Bio-Rad). For staining 3×10^6^ cells were plated in round bottom 96 well plates.

### Flow Cytometric Analysis

Single cells suspensions were incubated with anti-CD45 (1:100, Brilliant Violet 785, Biolegend), anti-CD11b (1:50, FITC, Biolegend) and live/dead (1:1000, eFlour 780 Fix Viability, Invitrogen) for 30 mins at 4°C. After incubation cold PBS was added to wash cells and samples were spun down at 1500 rpm for 5 minutes. For intracellular staining the Cytofix/Cytoperm kit (BD) was used according to manufacturer’s instructions. Briefly, fixation/permeabilization solution was added and cells were then incubated for 20 minutes at 4°C. Cells were then washed using perm/wash buffer to remove fixation solution. Cells were then stained with anti-iNOS (1:100, PE, Invitrogen) for 30 minutes at 4°C and then washed using perm/wash buffer. Single stained controls for compensation were done using anti-rat and anti-hamster Igκ/negative control beads (BD). Cell staining was analyzed using LSRFortessa X-20 cell analyzer (BD) at the CU | AMC ImmunoMicro Flow Cytometry Shared Resource. Flow analysis was then performed using FlowJo (BD).

### Statistical Analysis

Data analysis was performed with GraphPad PRISM. Data is shown as mean ± standard error of the mean (SEM) or min to max. One-way ANOVAs, Student’s t-tests, and Mann-Whitney U tests were performed. More information about statistical analysis can be found in individual figure legends.

## Acknowledgements

We would like to thank Andreas Baumler (University of California Davis) for providing the *S*. Typhimurium Kan^r^ and *S*. Typhimurium *narG narZ napA* Kan^r^ Carb^r^ strains (15). This work was funded by grants from the NIAID under Award Numbers AI164154 and AI173121, and the University of Colorado start-up funds to A.M.K.G.

## Author Contributions

L.A.S. and A.M.K.G wrote the manuscript. L.A.S. and A.M.K.G. designed experiments and L.A.S. and S.K.K.D performed the experiments. M.X.B. scored pathological changes. All authors commented on the manuscript.

