## Supplemental Table 1 for "Nitrate-mediated luminal expansion of *Salmonella* Typhimurium is dependent on the ER stress protein CHOP"

**Supplementary Table 1: Primers used in this study**

| Target | Forward Sequence (5’-3’) | Reverse Sequence (5’-3’) |
|---|---|---|
| *mGapdh* | TGTAGACCATGTAGTTGAGGTCA | AGGTCGGTGTGAACGGATTTG |
| *mHspa5* | GAGCGTCTGATTGGCGATGC | TTCCAAGTGCGTCCGATGAGG |
| *mXbp1* | GAGTCCGCAGCAGGTG | GTGTCAGAGTCCATGGGA |
| *mChop* | CTGGAAGCCTGGTATGAGGAT | CAGGGTCAAGAGTAGTGAAGGT |
| *mNos2* | TTGGGTCTTGTTCACTCCACGG | CCTCTTTCAGGTCACTTTGGTAGG |
| *mIl1b* | CCTGAACTCAACTGTGAAATGCC | TCTTTTGGGGTCCGTCAACTTC |
| *mKC* | TGCACCCAAACCGAAGTCAT | TTGTCAGAAGCCAGCGTTCAC |
| *mIl23* | CCAGCAGCTCTCTCGGAATC | TCATATGTCCCGCTGGTGC |
| *mTnfa* | AGCCAGGAGGGAGAACAGAAAC | CCAGTGAGTGAAAGGGACAGAACC |
