## Supplemental Figure Legends for "Nitrate-mediated luminal expansion of *Salmonella* Typhimurium is dependent on the ER stress protein CHOP"

**Supplemental Figures**

**Figure S1. Activation of the UPR in *Chop*$^{-/-}$ and control mice.** RNA from the colon of infected and uninfected *Chop*$^{-/-}$ and *Chop*$^{+/-}$ littermate control mice was extracted and analyzed by qRT-PCR to determine the levels of (A) Hspa5 and (B) Xbp1 mRNA expression at 24, 48 and 72hpi. (C and D) Streptomycin-pretreated *Chop*$^{+/+}$ and *Chop*$^{+/-}$ mice were infected with *S*. Typhimurium (SL1344, $10^8$ cfu/mouse) and bacterial numbers were determined in colon contents (A) and liver (B) at 24h, 48h and 72h post infection. Data shown as mean ± SEM or min to max with 4-9 mice per group. A and B; Multiple unpaired t tests. C and D; unpaired t tests, ns (not significant).

**Figure S2. Histopathology scores in *Chop*$^{-/-}$ and *Chop*$^{+/-}$ littermate control mice at 48hpi.** Streptomycin-pretreated *Chop*$^{-/-}$ and *Chop*$^{+/-}$ littermate control mice were infected with *S*. Typhimurium (SL1344, $10^8$ cfu/mouse) for 48h. (A) Total histopathology scores for colon sections (B) Total neutrophil count. Data shown as mean ± SEM with 5 mice per group. Unpaired t tests, ns (not significant).

**Figure S3. *Chop*$^{-/-}$ mice have reduced iNOS expression.** (A) Protein was extracted from colon sections of mice after 48 hours of *S*. Tm infection and Western blot analysis was used to determine the levels of iNOS and the loading control $\alpha/\beta$ tubulin. (B) Streptomycin-pretreated *Chop*$^{-/-}$ and *Chop*$^{+/-}$ littermate control mice were infected with a 1:1 mixture of wildtype *S*. Typhimurium SL1344 and *ΔnapAnarZnarG* ($10^8$ cfu/mouse). The competitive index (CI) in the colon contents was determined 72h post infection. Data shown as mean ± SEM with 9-10 mice per group. Unpaired t test, ns (not significant).

**Figure S4. *S*. Typhimurium-infected *Chop*$^{\Delta IEC}$ at 48hpi.** Streptomycin-pretreated *Chop*$^{\Delta IEC}$ and *Chop*$^{flox}$ littermate control mice were infected with *S*. Typhimurium (SL1344, $10^8$ cfu/mouse) and cfus were determined in colon contents (A) and liver (B) at 72hpi. (C) Total histopathology scores and (D) neutrophil counts in the colon at 48hpi. (E-I) *Xbp1, Hspa5, Nos2, Kc and Il1b* mRNA levels in the colon. Data shown as mean ± SEM or min to max with 4-6 mice per group. (A and B) Mann-Whitney U test, (C-I) unpaired t tests, ns (not significant).

**Figure S5. Cytokine expression in *S*. Typhimurium-infected *Chop*$^{\Delta IEC}$ at 72hpi.** RNA from the colon of infected and uninfected *Chop*$^{\Delta IEC}$ and *Chop*$^{flox}$ littermate control mice was extracted and analyzed by qRT-PCR to determine the levels of *Kc* (A), *Il1b* (B), *Tnfa* (C) and *Il23* (D) at 72h post infection. Data shown as mean ± SEM with 5 mice per group. Unpaired t tests, ns (not significant).
