## Supplementary figures and images for "Nitrate-mediated luminal expansion of *Salmonella* Typhimurium is dependent on the ER stress protein CHOP"

### Supplemental Figure 1

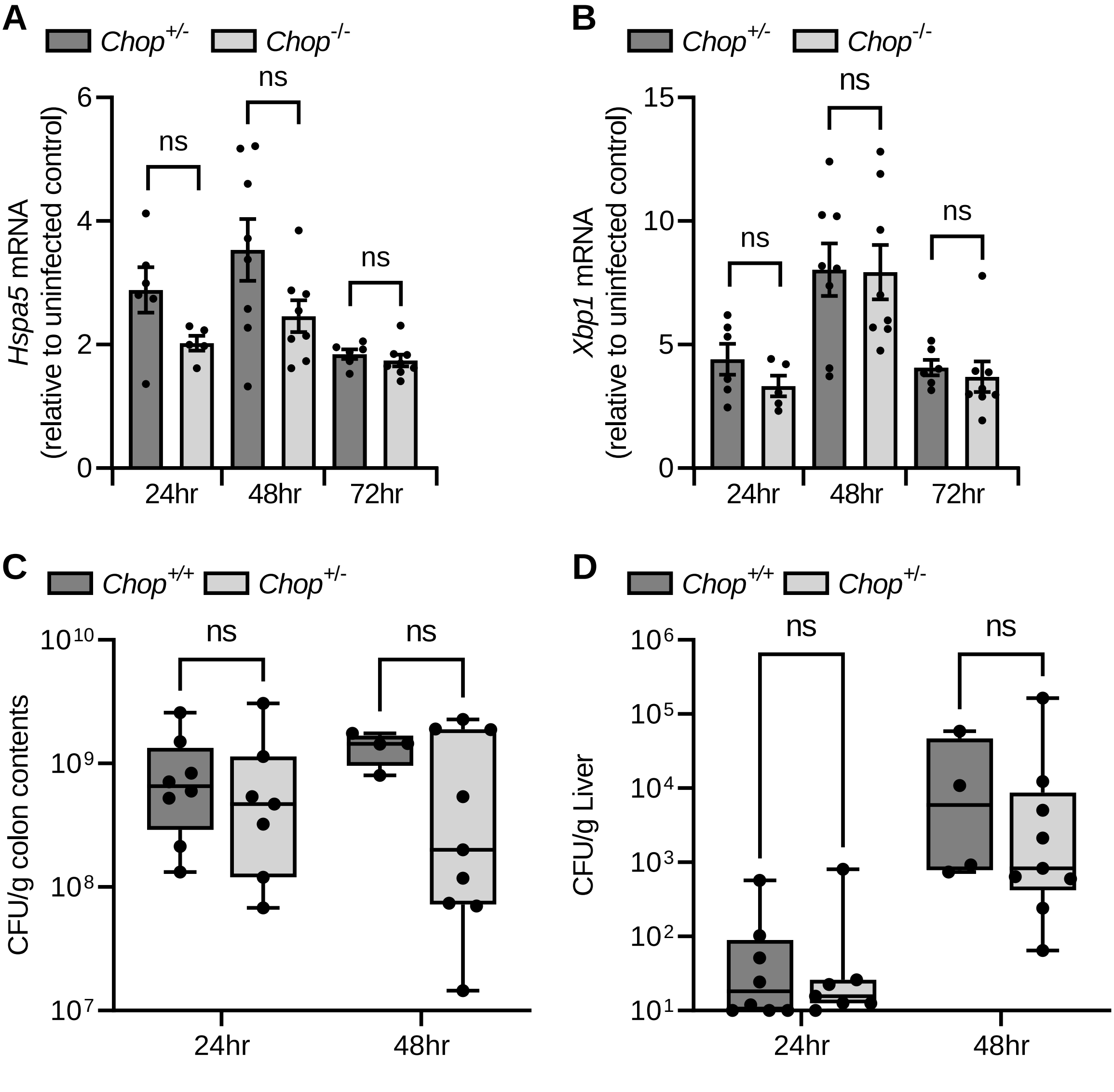

### Supplemental Figure 2

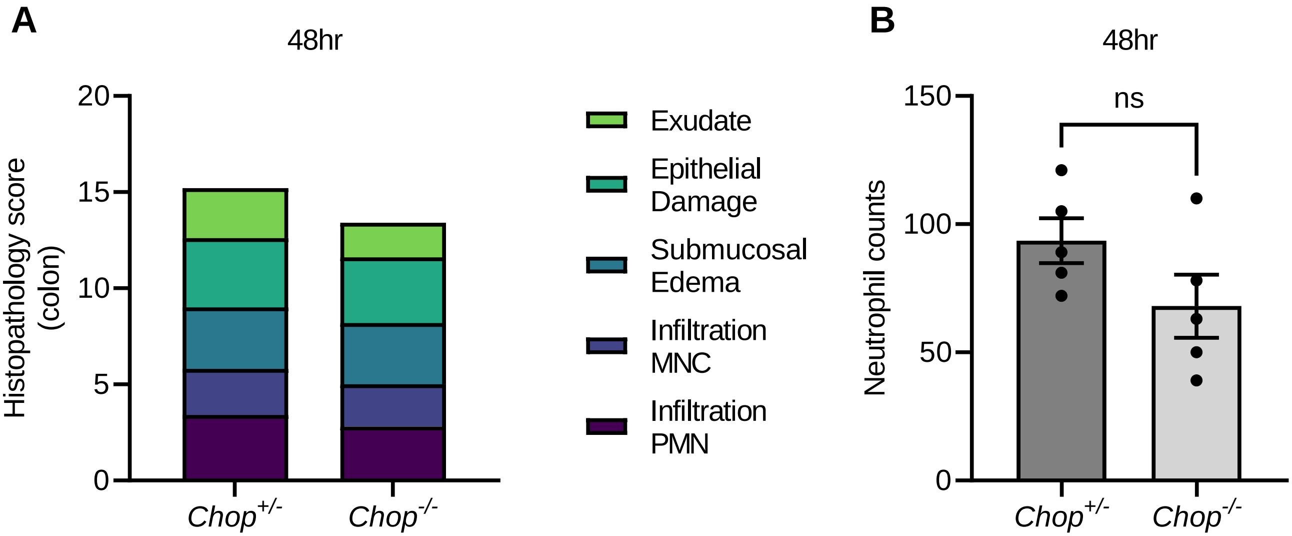

### Supplemental Figure 3

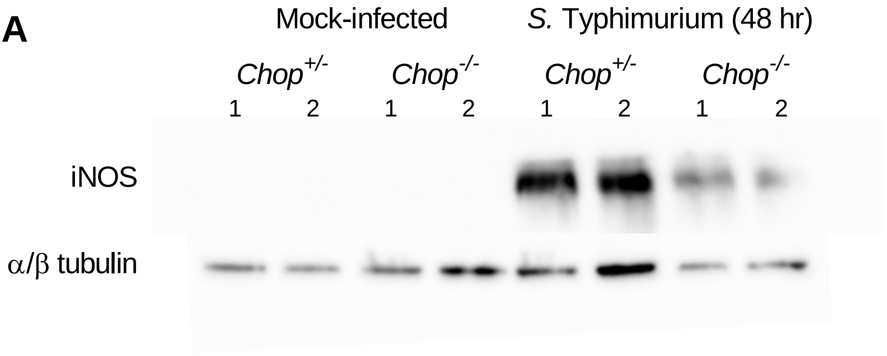

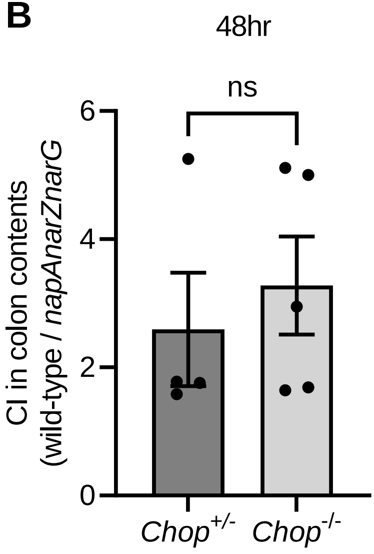

### Supplemental Figure 4

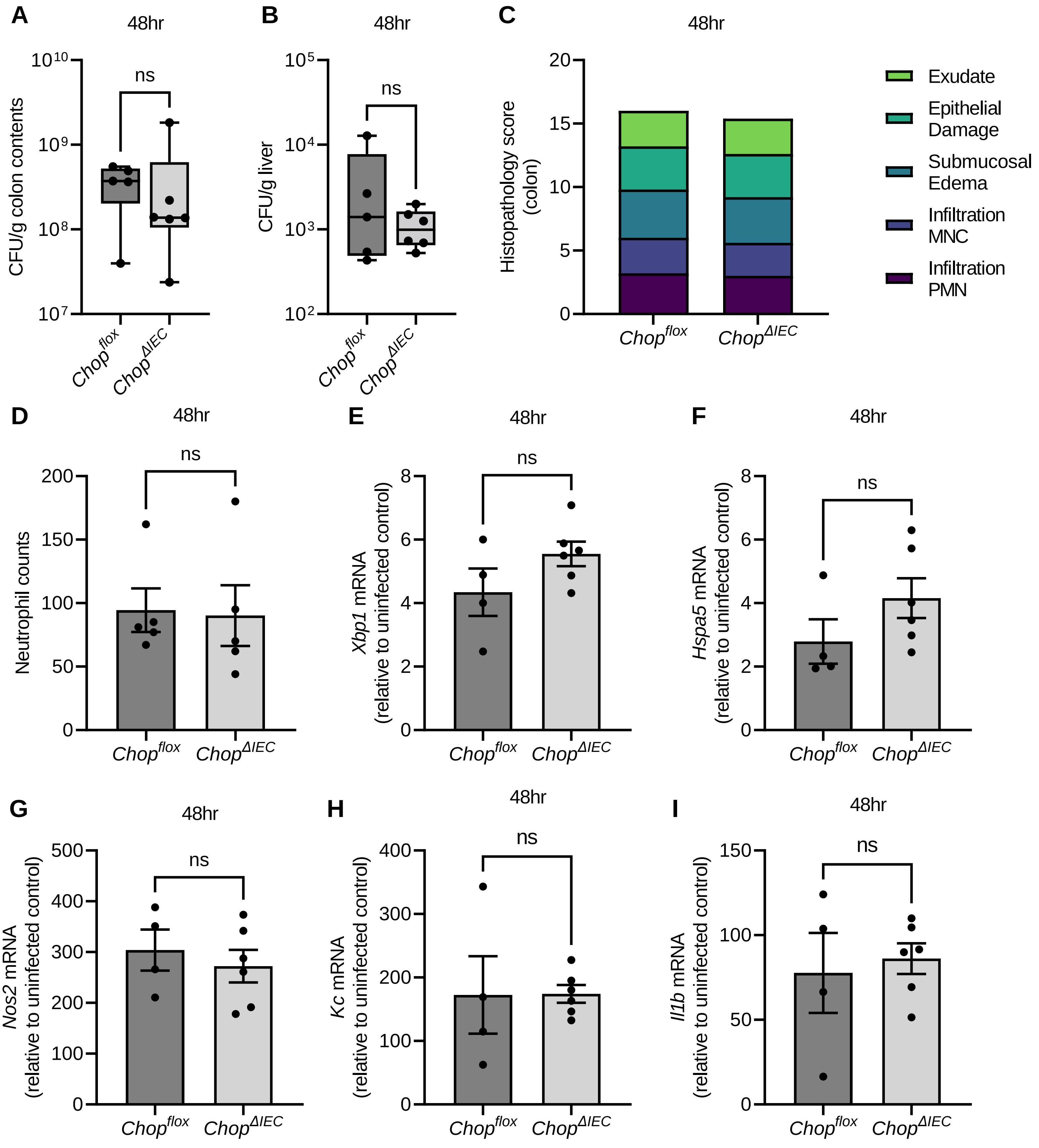

### Supplemental Figure 5

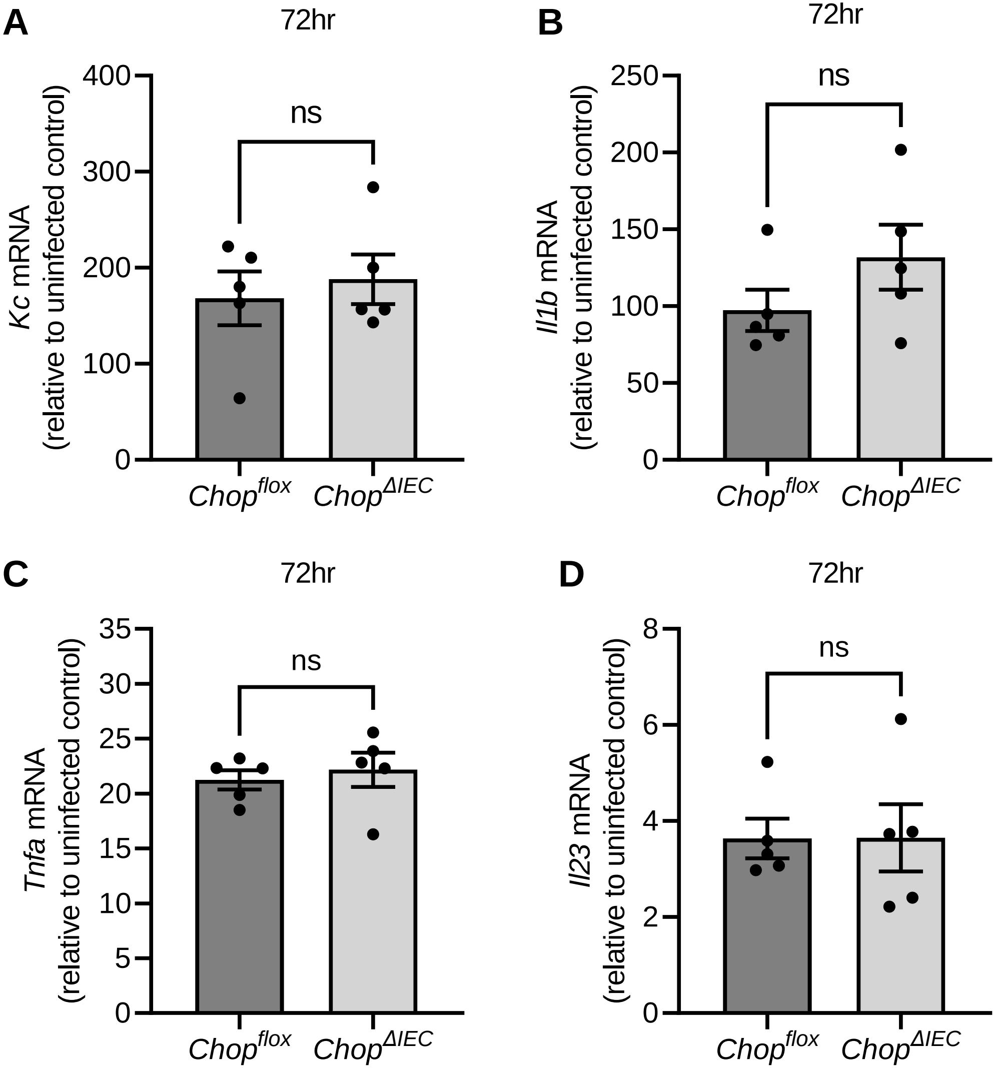
